# Multimodal brain age prediction using machine learning: combining structural MRI and 5-HT2AR PET derived features

**DOI:** 10.1101/2024.02.05.578968

**Authors:** Ruben P. Dörfel, Joan M. Arenas-Gomez, Claus Svarer, Melanie Ganz, Gitte M. Knudsen, Jonas E. Svensson, Pontus Plavén-Sigray

## Abstract

To better assess the pathology of neurodegenerative disorders and the efficacy of neuroprotective interventions, it is necessary to develop biomarkers that can accurately capture age-related biological changes in the human brain. Brain serotonin 2A receptors (5-HT2AR) show a particularly profound age-related decline and are also reduced in neurodegenerative disorders, such as Alzheimer’s disease.

This study investigates whether the decline in 5-HT2AR binding, measured in vivo using positron emission tomography (PET), can be used as a biomarker for brain aging. Specifically, we aim to 1) predict brain age using 5-HT2AR binding outcomes, 2) compare 5-HT2AR-based predictions of brain age to predictions based on gray matter (GM) volume, as determined with structural magnetic resonance imaging (MRI), and 3) investigate whether combining 5-HT2AR and GM volume data improves prediction.

We used PET and MR images from 209 healthy individuals aged between 18 and 85 years (mean=38, std=18), and estimated 5-HT2AR binding and GM volume for 14 cortical and subcortical regions. Different machine learning algorithms were applied to predict chronological age based on 5-HT2AR binding, GM volume, and the combined measures. The mean absolute error (MAE) and a cross-validation approach were used for evaluation and model comparison.

We find that both the cerebral 5-HT2AR binding (mean MAE=6.63 years, std=0.74 years) and GM volume (mean MAE=6.95 years, std=0.83 years) predict chronological age accurately. Combining the two measures improves the prediction further (mean MAE=5.54 years, std=0.68). In conclusion, 5-HT2AR binding measured using PET might be useful for improving the quantification of a biomarker for brain aging.

## Introduction

Aging is associated with a wide range of molecular and cellular changes throughout an individual’s lifespan, resulting in the deterioration of physical condition [1–3]. Biological changes caused by aging are the main drivers for overall mortality and chronic pathology [4]. Pathological brain aging results in cognitive impairment [5] and predisposes individuals for neurodegenerative disorders, such as Alzheimer’s and Parkinson’s disease [6]. To describe biological changes preceding the onset of neurodegenerative disorders, and to assess the effects of interventions meant to alleviate the effect of biological aging, it is necessary to develop biomarkers that can robustly capture such changes [2,7].

By using large neuroimaging datasets covering the human lifespan, it is possible to study changes in aging-related brain biology noninvasively. Structural magnetic resonance imaging (MRI) studies have shown a robust age-related decrease in gray matter (GM) and white matter volumes, with different rates of decline over the lifespan [8]. Most work so far has been done using structural MRI, where large datasets are publicly available, but other MR protocols and neuroimaging modalities have also been used to study the effects of aging on the brain. Functional MRI data supports a decline in functional connectivity [9], and diffusion-weighted MRI studies have linked demyelination and axonal degeneration to increasing age [10]. Positron emission tomography (PET) allows for in vivo quantification of proteins and processes that change with increasing age, such as metabolism, neuroinflammation, and many neurotransmitter systems [11–13].

A common observation in many different PET tracers is an association between radioligand binding and age [14,15]. One receptor type that shows a particularly strong age-related decline in PET studies is the serotonin 2A receptor (5-HT2AR) [12,16–21]. In the human brain, 5-HT2ARs are mainly located in the neocortex layer V and, to a lesser degree in subcortical regions such as the hippocampus and putamen [14]. The 5-HT2AR is reduced in a number of neurodegenerative diseases, such as Alzheimer’s disease [22–24].

Machine learning (ML) allows for modeling the healthy aging brain by summarizing aging brain biology from neuroimages into one single variable, called “*brain age”* [25,26]. Models are trained to predict chronological age on imaging data from healthy individuals based on structural or functional change that occur over the human lifespan. The result is a model of a healthy aging brain, indicating how an average healthy brain would change during an individual’s lifespan. More than the predicted age itself, the deviation of the predicted age from the true chronological is of interest. This deviation is hypothesized to reflect divergence from the expected healthy aging trajectory [27]. The deviation between predicted age (Â) and chronological age (*A*) is expressed as the predicted age deviation (PAD):

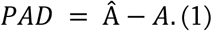

Under this hypothesis, a higher predicted than chronological age reflects a biologically older brain. Following from this, a positive PAD should translate to an increased risk for aging-related disorders. An increased PAD has been linked to increased cardiovascular risk [28], overall mortality [27], neurodegenerative disorders, and cognitive decline [29–33].

Here we aim to investigate if the age-related decline in 5-HT2AR binding can be utilized in the brain age paradigm. Specifically, the first aim is to apply ML algorithms to 5-HT2AR binding outcomes from PET images to predict brain age. The second aim is to compare these estimates to those derived from applying the same algorithms to volumetric GM data derived from structural MR images. The third aim is to investigate whether a multimodal approach combining 5-HT2AR binding and GM measures improves the prediction of brain age.

## Materials and Methods

### Imaging Data

#### Dataset

The Center for Integrated Molecular Brain Imaging (Cimbi) database [34] stores a large collection of high-resolution 5-HT neuroimaging data from healthy individuals. Here we included healthy participants from the Cimbi database where 5-HT2AR PET and structural MRI data were available. In total 209 individuals were included in the study. For imaging, the two PET radioligands [^11^C]Cimbi-36 [35] or [^18^F]altanserin [36] were used. The demographics and PET acquisition details are summarized in Table 1.

**Table 1:** The demographics of the used subsets.

|  | Scans | Gender<br>Male / Female | Age<br>min/max | Age<br>mean (std) | PET Camera<br>GE / HRRT | MR Scanner<br>1.5T / 3T |
| --- | --- | --- | --- | --- | --- | --- |
| [<sup>11</sup>C]Cimbi-36 | 67 | 39 / 28 | 18.4 / 58.3 | 28.3 (7.7) | - / 67 | - / 67 |
| [<sup>18</sup>F]altanserin | 142 | 90 / 52 | 18.5 / 81.7 | 40.5 (18.5) | 142 / - | 70 / 72 |
| Total | 209 | 129 / 80 | 18.4 / 81.7 | 36.6 (16.8) | 142 / 67 | 70 / 139 |

#### PET and MRI image acquisition

The MRI scans were acquired on Siemens (Siemens, Erlangen, Germany) scanners with a field strength of either 1.5T or 3T using 3D magnetization-prepared rapid gradient-echo sequences. All [^11^C]Cimbi-36 PET scans were acquired on a Siemens High-resolution Research Tomograph system (HRRT; CTI/Siemens, Knoxville, TN, USA), whereas all [^18^F]altanserin scans were acquired on a GE-Advance scanner (GE, Milwaukee, WI, USA). [^11^C]Cimbi-36 was administered as an intravenous bolus injection with data collected for two hours [35]. [^18^F]altanserin was administered intravenously, starting with a bolus injection followed by continuous infusion. Data acquisition started two hours after [^18^F]altanserin injection to allow for a steady state to be reached and lasted for 40 minutes. For all [^18^F]altanserin scans, a venous blood sample was obtained, and metabolite-corrected radioactivity was quantified to allow for the correction of radionuclide-labeled metabolites entering the brain [37].

#### Image pre-processing

All PET images were corrected for inter-frame motion using the AIR algorithm (v5.3.0) [38]. The MR images have been corrected for spatial distortions due to nonlinearities in gradient fields. This preprocessing was part of the Cimbi database curation.

Standard FreeSurfer (v7.2) segmentation and parcellation pipelines [39] were applied to the MR images to retrieve masks for cortical and subcortical regions of interest (ROIs) from the Desikan-Killiany [40], and Aseg atlas [41]. Cortical ROIs were grouped into seven larger cortical regions (frontal lobe, occipital lobe, parietal lobe, temporal lobe, parahippocampal gyrus, cingulate, and insula) [42] with high 5-HT2AR binding, as identified by autoradiography [14,43]. Additionally, seven subcortical brain regions were used in the analysis (thalamus, caudate, putamen, pallidum, hippocampus, amygdala, and nucleus accumbens). All regions were averaged across hemispheres, resulting in 14 ROIs (Table S1). The MR images were co-registered to a summed PET image using SPM8. The resulting co-registration matrix was used to project the ROI masks on the dynamic PET files to extract time-activity curves (TACs) for each region.

### Brain Age Modelling

#### Outcome Measures

The volume of all ROIs was extracted from the FreeSurfer output and normalized using the intracranial volume [44]. Regional binding potentials (BPs) were calculated using the regional TACs extracted from the PET data. The cerebellum excluding vermis was used as a reference region, as it has a negligible density of 5-HT2ARs [35]. 5-HT2AR binding from [^11^C]Cimbi-36 scans was quantified as the non-displaceable binding potential (*BP*_*ND*_) using multilinear reference tissue model 2 [45], with k2’ values estimated using the simplified reference tissue model applied to a TAC extracted from a mask over the entire neocortex. Quantification of [^18^F]altanserin binding has been described in detail elsewhere (Pinborg et al., 2003). In short, the ratio of specifically bound radioligand to that of total parent radioligand in plasma, (*BP*_*P*_) was calculated and used as the outcome measure. Even though *BP*_*P*_ and *BP*_*ND*_ are different measures, both are proportional to the density of the 5-HT2AR and highly correlated [46].

#### Machine learning algorithms

To develop a model of the healthy aging brain, a set of ML algorithms were trained to predict chronological age from imaging-derived GM volume and 5-HT2AR binding data. The algorithms were implemented using scikit-learn (v1.2) [47] in python 3.9.12. We selected commonly used algorithms for brain age prediction [48,49]: 1) Bayesian Ridge Regression (BRidge), 2) Relevance Vector Regression (RVR) implemented using *ARDRegression*, 3) Gaussian Process Regression with linear kernel (linGPR) 4) Gaussian Process Regression with radial basis function kernel (rbfGPR), and 5) linear support vector regression (linSVR). The selected algorithms were trained on either 5-HT2AR binding outcomes, GM volumes, or both. The multimodal model, combining structural MRI and 5-HT2AR PET-derived data, was implemented as a stacking regressor [50]. In a stacking regressor, a base model for each modality was trained, with outcomes then used as input into a linear regression model. The algorithms BRidge, RVR, linGPR, linSVR, and rbfGPR were used as a base estimator, resulting in five ensemble regressors.

#### Baseline predictors

Two references were implemented to put the results into perspective. First, we trained a dummy regressor that always outputs the mean age of the training. Second, we applied pyment, a pre-trained state-of-the-art structural MRI-based brain age prediction software. pyment uses a skull-stripped and MNI-space registered T1w-image as input [51]. It was originally trained on images from 53542 healthy individuals and has been shown to predict age on unseen data with high accuracy and reliability [52].

#### Experiments

We predicted chronological age based on the three feature sets (5-HT2AR, GM, and 5-HT2AR + GM) as well as the two reference models. An overview of the respective prediction pipelines is presented in Figure 1. The different outcome measures for [^11^C]Cimbi-36 and [^18^F]altanserin were aligned by transforming regional 5-HT2AR *BP*_*ND*_ to *BP*_*P*_-like values based on age-matched subsets using a distribution mapping approach [53], described in the supplement (S1.2). Subsequently, the matched values were standardized (zero mean, unit variance). GM volumes were similarly z-scored. A multimodal brain age estimate was calculated combining 5-HT2AR binding and GM volume features using the stacking approach previously described. Each feature set was transformed as described before.

**Figure 1:**
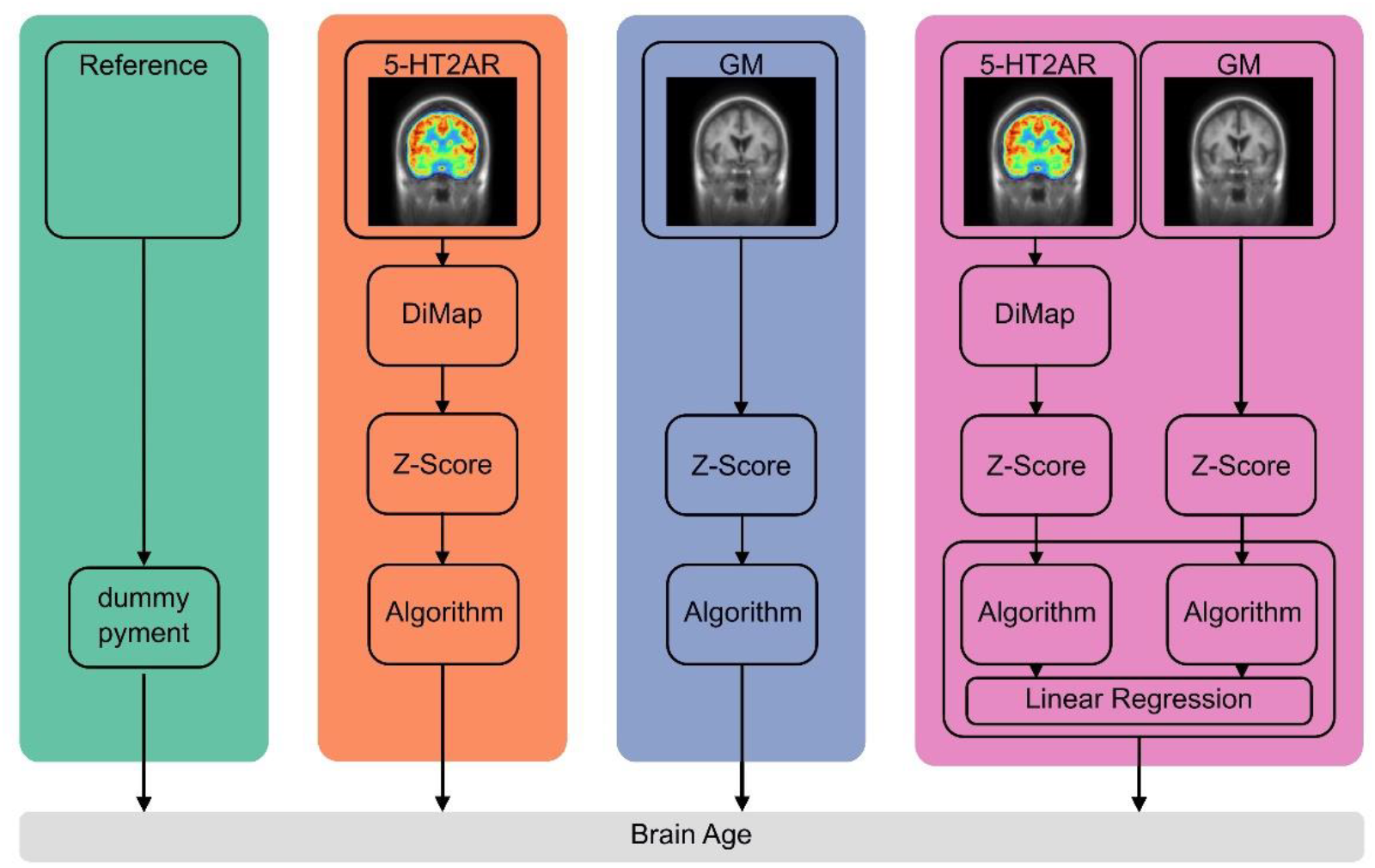
Overview of the prediction pipelines. **Reference:** the pre-trained MRI-based prediction model pyment and a dummy regressor always predicting the mean chronological age training data. **5-HT2AR:** *BPND* ([^11^C]Cimbi-36) is transformed to *BPP* -like units using the distribution mapping (DiMap). BPs are then z-scored. **GM:** Volumes of cortical and subcortical regions are z-scored. **5-HT2AR+GM:** Transformations for 5-HT2AR and GM features are identical to the unimodal counterparts. Brain age is predicted using a stacking ensemble method where a base model is trained for each modality, with the outcome used as input into a final estimator.

#### Model training and evaluation

Model accuracy was assessed using the mean absolute error (MAE), calculated as the average absolute difference between predicted and chronological age for each validation fold. The generalization error of each model was estimated based on the 100 hold-out folds in a 20-times repeated 5-fold cross-validation (CV) setup [54]. An additional inner 5-fold CV was used for hyperparameter tuning, if necessary. The initial split was randomly shuffled for each of the 20 repetitions to gain a split-independent estimate of the generalization error. The outer five-fold CV was stratified by age to ensure the same age distribution between the training and validation sets.

The generalization error for each model was then estimated as the average MAE over all folds (N=100)

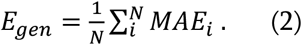

Further, Pearson’s *r* was calculated between chronological and brain age. The correlation coefficient should be close to one if the estimated brain age perfectly follows the chronological age in this population of healthy participants. Pearson’s r was also calculated between chronological age and PAD (see eq. 1) values. Here, the correlation coefficient should be close to zero as the PAD is supposed to reflect deviations from the healthy aging trajectory and should not be associated with chronological age. The coefficient of determination (*R*^2^) was calculated to assess how much variance in the predicted variable (chronological age) was explained by the respective model. To describe the strength of the association or explained variance, we used the following heuristic: an *r/R2* between 0–0.19 was described as negligible, 0.2–0.39 as weak, 0.40–0.59 as moderate, 0.6–0.79 as strong/large and 0.8–1 as very strong/large [55].

#### Model Comparisons

To assess whether GM volumes are more predictive of age than 5-HT2AR binding in the same regions and whether a multimodal approach (combining 5-HT2AR+GM features) can improve predictions, we tested the best model trained on GM or 5-HT2AR+GM features against the best model trained on only 5-HT2AR features using a paired two-tailed t-test. Specifically, the MAE per fold for each model was used for comparison. As most of the training data overlaps in each CV split, the assumption of independence between the CV folds within each repetition was violated. Therefore, a correction to the t-test was applied to adjust for this dependence [56,57]. The significance level was set to 0.05.

#### Modality contribution and feature importance

To investigate the similarity between predictions, we correlated predictions of the models based on GM volumes and 5-HT2AR binding. A high correlation coefficient would indicate that the information in both feature sets leads to similar brain age predictions. As an underlying strong correlation with chronological age is expected, we also correlated the PAD values from the different feature sets. Additionally, we investigated the weight for each modality in the stacking process of the combined model. This indicates which feature set contributes most to the final predictions of the combined model.

The BRidge algorithm allows for easy retrieval and interpretation of fitted parameters. For this reason, we used the model BRidge for this analysis. The model weights were obtained and averaged over CV folds for this analysis. These average values were then displayed for each ROI on a standard T1w template image. The weights indicate how important a feature was, i.e., how much it contributed to the final prediction in combination with the other features. This analysis was performed post-hoc.

## Results

### Model evaluation and comparison

Brain age was predicted using two reference models (dummy, pyment) and a set of models trained on three different feature sets: 5-HT2AR binding, GM volumes, and the combination of 5-HT2AR + GM data. The results for the best-performing model in each feature set are visualized in Figure 2 and summarized in Table 2. A complete comparison of all models can be found in supplement S2.1. Overall, all models performed better than the dummy regressor and worse than the state-of-the-art software pyment. Figure 3 shows the mean prediction for each subject over the 20 iterations on each feature set for the specific best performing model. There was a strong correlation between chronological and predicted age and a moderate to strong negative correlation between PAD and chronological age.

**Table 2:**
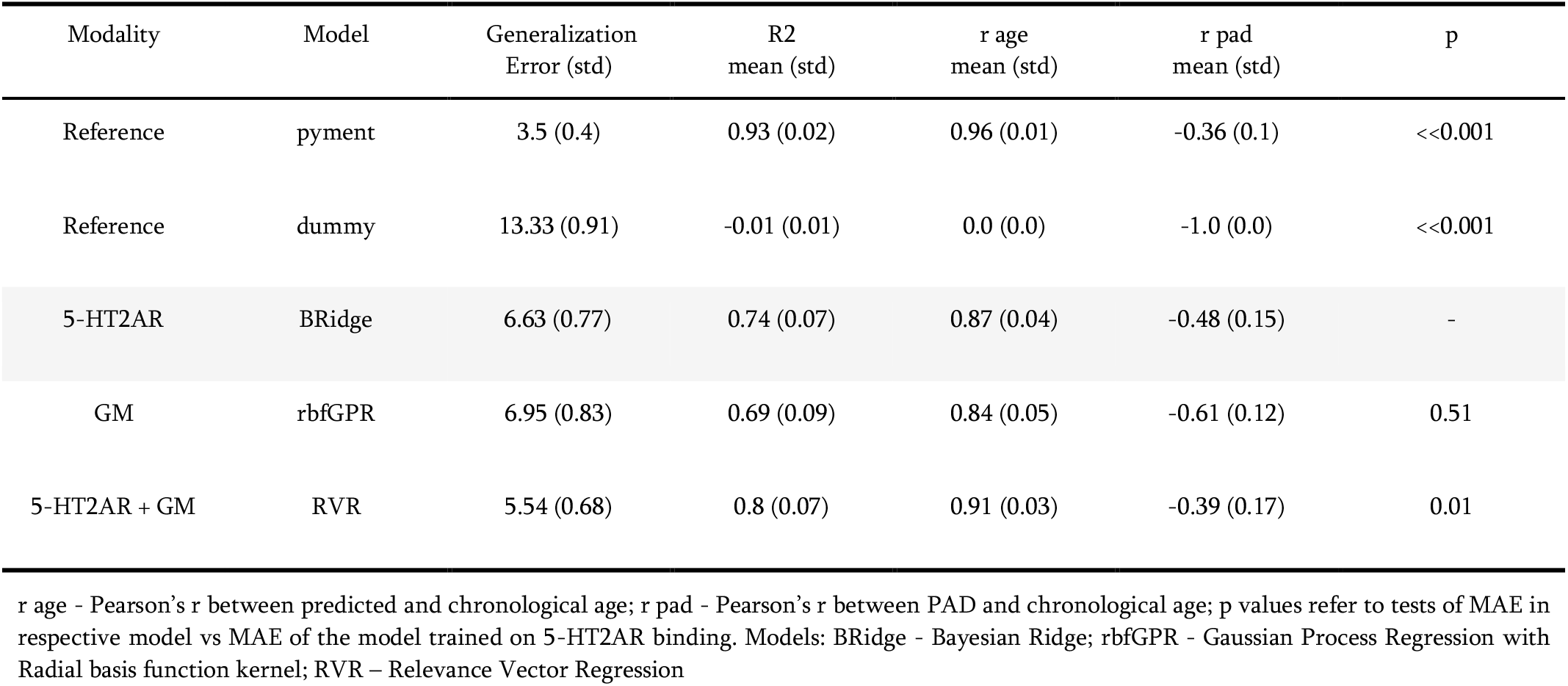
The results of the best-performing model for the four experiments (Reference, 5-HT2AR, GM, 5-HT2AR + GM). Values are mean (std) over the 100 CV iterations for MAE, R2, r age, and r pad.

**Figure 2:**
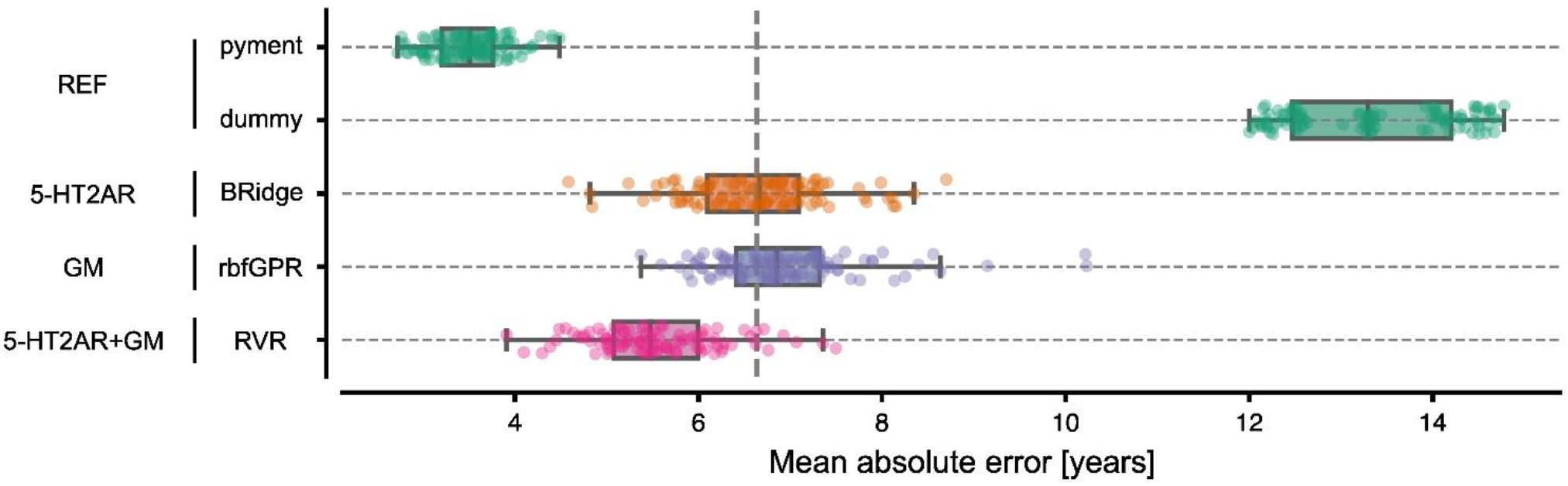
MAE for the best-performing models for each feature set. Each dot represents the MAE for one of the 100 CV iterations.

**Figure 3:**
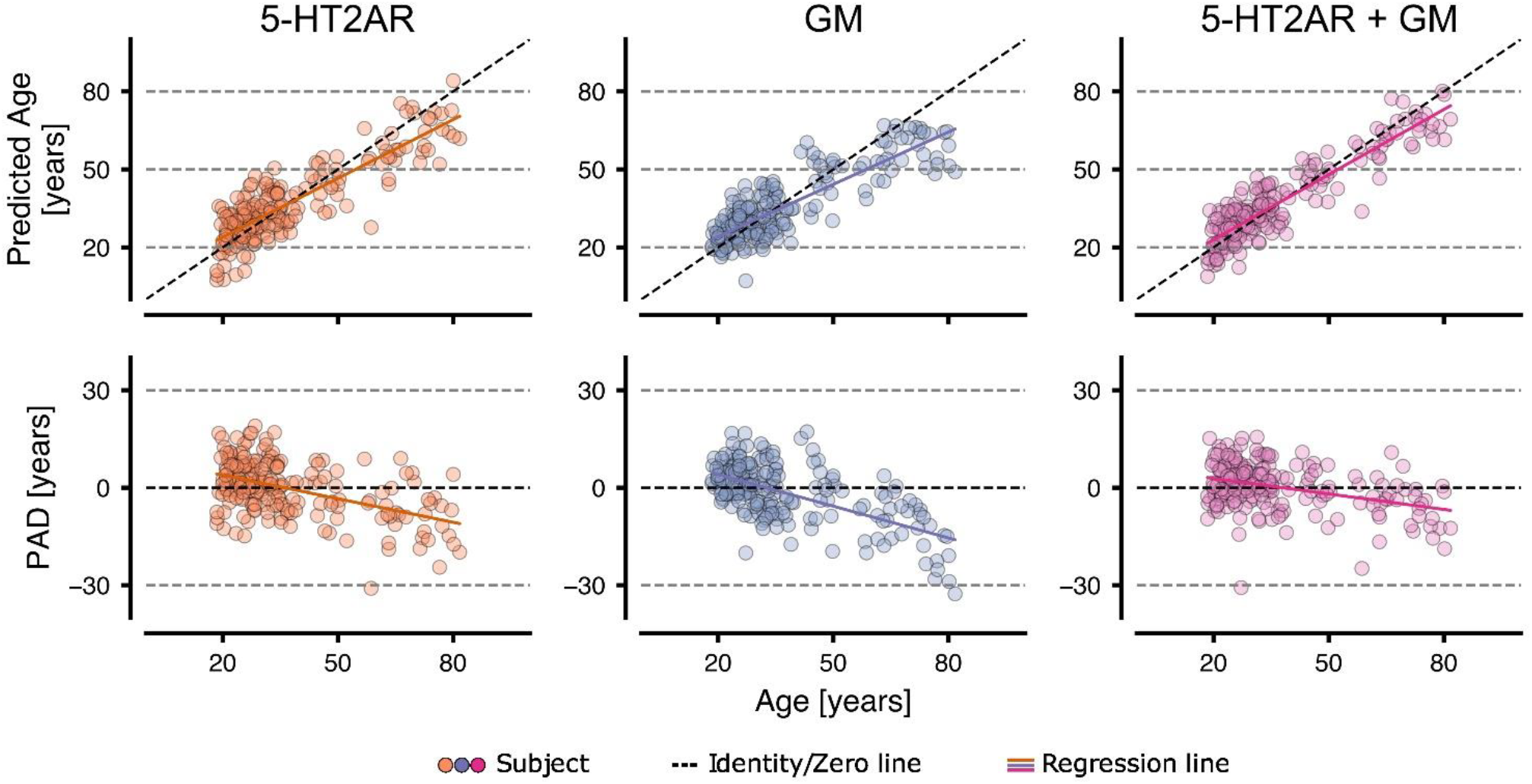
The predicted age and PAD vs. the chronological age for 5-HT2AR, GM, and 5-HT2AR+GM using the best performing model (5HT2AR - Bridge; GM - rbfGPR; 5HT2AR+GM - RVR). Each point represents a mean value over 100 CV iterations. The colored line represents the fitted regression line.

The 5-HT2AR-based model showed that chronological age in healthy individuals could be predicted using regional 5-HT2AR binding with a generalization error of 6.63 years and a very strong correlation between predicted and chronological age (r=0.87). A moderate negative correlation between PAD and age (r=-0.48) was observed.

The GM-based model showed that chronological age in healthy individuals could be predicted using cortical and subcortical GM volumes with a generalization error of 6.95 years and a very strong correlation (r=0.84) between predicted and chronological age. Further, a strong negative correlation between PAD and age (r=-0.61) was observed. The predictions were not significantly different than those based on 5-HT2AR binding (p = 0.51).

The combined 5-HT2AR+GM-based model showed that chronological age in healthy individuals could be predicted using a combination of regional 5-HT2AR binding potential and cortical and subcortical GM volumes, with a generalization error of 5.54 years and a very strong correlation (r=0.91) between predicted and chronological age. A negligible negative correlation between PAD and age (r=-0.39) was observed. The predictions were more accurate than those from both unimodal models. The statistical test comparing the 5-HT2AR-based model with the combined model was statistically significant (p = 0.01).

### Feature importance and modality contribution

We also investigated the contributions of each modality and ROI to the brain age predictions. For the sake of clarity, the results for the Bayesian Ridge Regressor were used in this analysis, since it is a linear model and weights are easily derived. The correlation between the PAD from the models based on 5-HT2AR binding vs. GM volume feature sets was r=0.35 with R2=0.12 (Figure 4A). Figure 4B shows the weight of each feature set in the ensemble regressor, with 5-HT2AR-based predictions contributing more to the final outcome. The contribution from individual ROIs to the predicted age for each base model is visualized in Figure 4C and reported as boxplots in the supplement (Figure S2.2). For 5-HT2AR binding, the temporal and parietal lobes, insula, caudate, and putamen displayed the highest weights. The frontal lobe and insula contributed the most to age prediction based on GM volumes.

**Figure 4:**
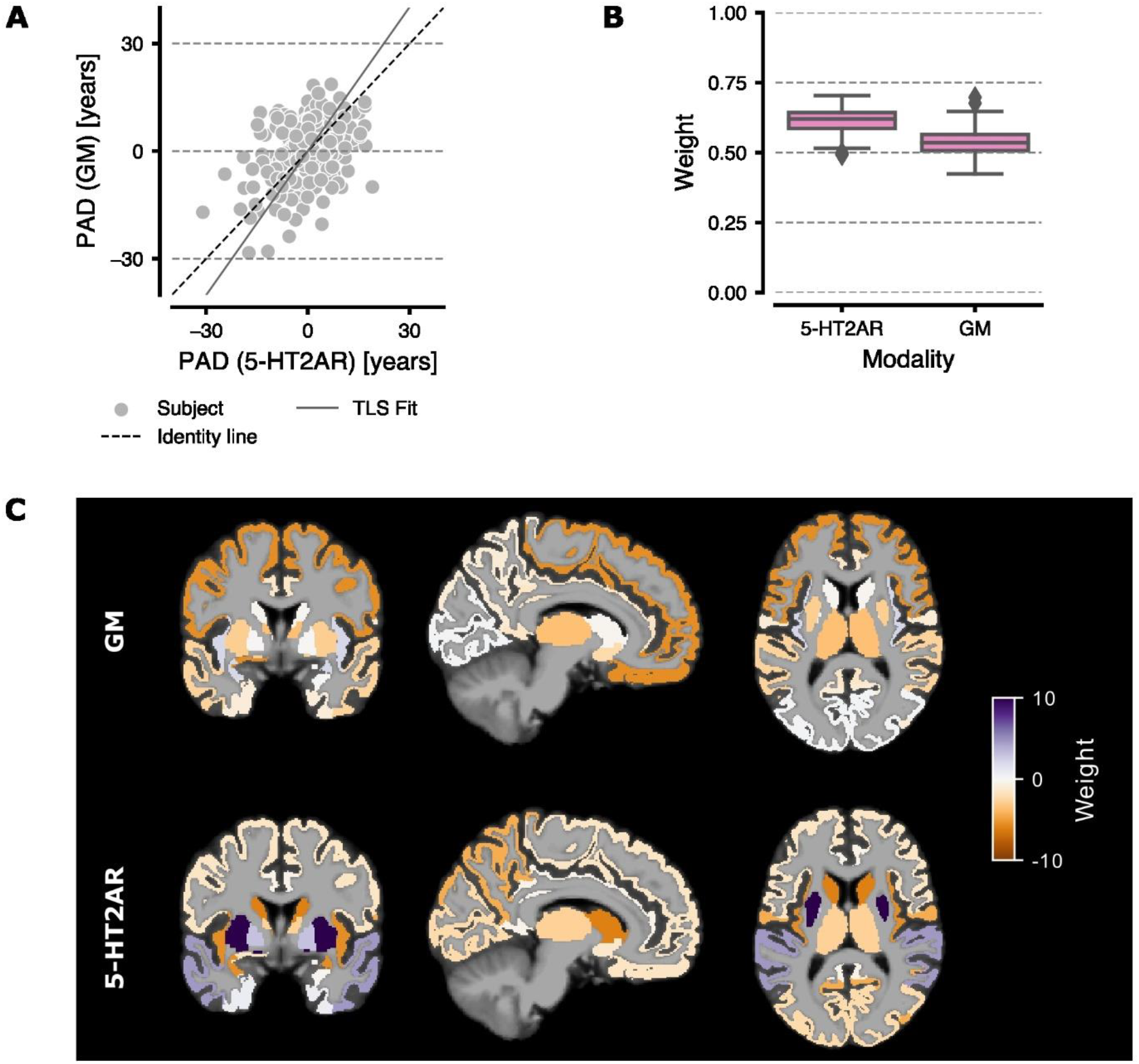
Comparison of 5-HT2AR vs GM-based models. Panel **A)** The PAD from GM volume-based predictions vs 5-HT2AR binding predictions. The solid gray line marks the regression line fitted using total least squares (TLS). Each point represents the mean PAD per subject averaged over all 100 CV iterations. Panel **B)** Boxplots showing the weights for the final ensemble regressor. Whiskers show 1.5 the interquartile range. Panel **C)** The weights for the ensemble base regressors (BRidge) overlaid on a MNI152 template brain.

## Discussion

This study investigated whether 5-HT2AR binding can be used as a putative biomarker for brain aging. For this purpose, we trained ML algorithms to predict chronological age based on regional binding estimates from 5-HT2AR PET images. We also trained ML models to predict chronological age from MRI GM volume estimates from the same regions, and we investigated whether combining both feature sets could improve overall performance. We found that ML algorithms trained on regional 5-HT2AR binding predicted chronological age with similar accuracy to those using regional GM volume as input. Further, combining data from both feature sets significantly improved the overall performance compared to both unimodal models.

The decline in 5-HT2AR binding with age is well described in the literature [12]. However, it is unclear to what extent the decline in binding estimates is caused by aging-related biology affecting the serotonin system or by structural changes affecting the PET signal. As the gray matter volume in most regions of the human brain decrease with age [8], an increased “spill-out” of radioactivity will be registered by the PET system due to a limited image resolution [58]. This could lead to lower observed binding estimates in brain regions while the true concentration of the 5-HT2AR is unchanged. In our data, age predictions based on 5-HT2AR binding performed similarly to age predictions based on GM volumes. However, combining the two feature sets showed increased accuracy over both unimodal approaches, indicating that the decline in 5-HT2AR binding contains unique information when predicting age, beyond that of volumetric changes in the brain. Further, figure 4C shows that different regions contribute to the respective age predictions for each feature set, indicating different aging patterns. This supports the notion that the decline in 5-HT2AR binding is unlikely to be caused solely by radioactivity spill-out.

A significant correlation between PAD and chronological age was observed for all models. Such a correlation is a well-known issue discussed extensively in the brain age literature [59]. Generally, a less accurate model predicting chronological age also shows a stronger negative correlation between PAD and chronological age. Such an association is problematic since the PAD is supposed to reflect a higher (or lower) biological age and should not mirror any variance in chronological age.

The predictions from the pre-trained MRI based model pyment showed high accuracy and outperformed all other tested methods. This finding was expected, since pyment predicts age based on voxels from the entire brain and is pre-trained on an extensive training dataset (> 50,000 T1w MRI scans). Due to limitations in the sample size used in the current study (N=209), we decided to use regional (ROI) level data. Since white matter and the ventricle volumes change with age [8], it is likely that including such features would have improved the predictions for the MRI-based models in our study. It should however be noted that the observed results from both the 5-HT2AR binding and GM volume based models were on par with the accuracy of other brain age prediction models in the literature [48,49,60]. The improved accuracy of predictions based on 5-HT2AR binding over predictions based on GM volumes indicates that with increased sample size for training, multi-modal brain age estimation incorporating 5-HT2AR features might outperform state-of-the-art MRI-based methods.

Our results are based on data from healthy individuals, conclusions from the current study are limited to this healthy population. However, previous PET studies have shown that, compared to age-matched healthy controls, cerebral 5-HT2AR binding is reduced in mild cognitive impairment and in Alzheimer’s disease [22,61]. This suggests that a brain age model based on 5-HT2AR binding could be useful in predicting neurodegeneration. Future studies should investigate whether 5-HT2AR based brain age estimation is useful as a biomarker for pathological aging and can be used for e.g., early detection of neurodegenerative disorders or for evaluating putative neuroprotective interventions intended to slow age-related processes in the human brain.

## Conclusion

We find that cerebral 5-HT2AR binding across different brain regions predicts chronological age with similar accuracy as MR-based gray matter measurements. Combining both measures significantly improved the prediction of age, indicating that both contribute unique information about brain aging to the model. We propose combined machine learning models of 5-HT2AR binding and GM volume estimates as an effective biomarker for aging- or disease-related changes in the human brain.

## Acknowledgements

We would like to thank Kristoffer Hougaard Madsen for his valuable comments in his capacity as censor for the master thesis preceding this manuscript.

## Notes

**Funding:** This work was supported by a Longevity Impetus Grant from the Norm Group. JS was supported by a postdoctoral grant from The Swedish Brain Foundation. PPS was supported by the Swedish Research Council (grant 2021-00462).

**Competing Interests:** The authors declare no conflicts of interest in relation to this work.

### Competing Interest Statement

The authors have declared no competing interest.

### Summary of Updates

Internal reviewers alerted us to a problem with FreeSurfer's estimation of the intracranial volume (ICV), which was used throughout our analysis. Consequently, we recalculated the ICV using SPM and updated the relevant sections accordingly.

