## Supplemental Information for "Multimodal brain age prediction using machine learning: combining structural MRI and 5-HT2AR PET derived features"

### Supplement

#### S1 Methods

##### S1.1 Regions of Interest

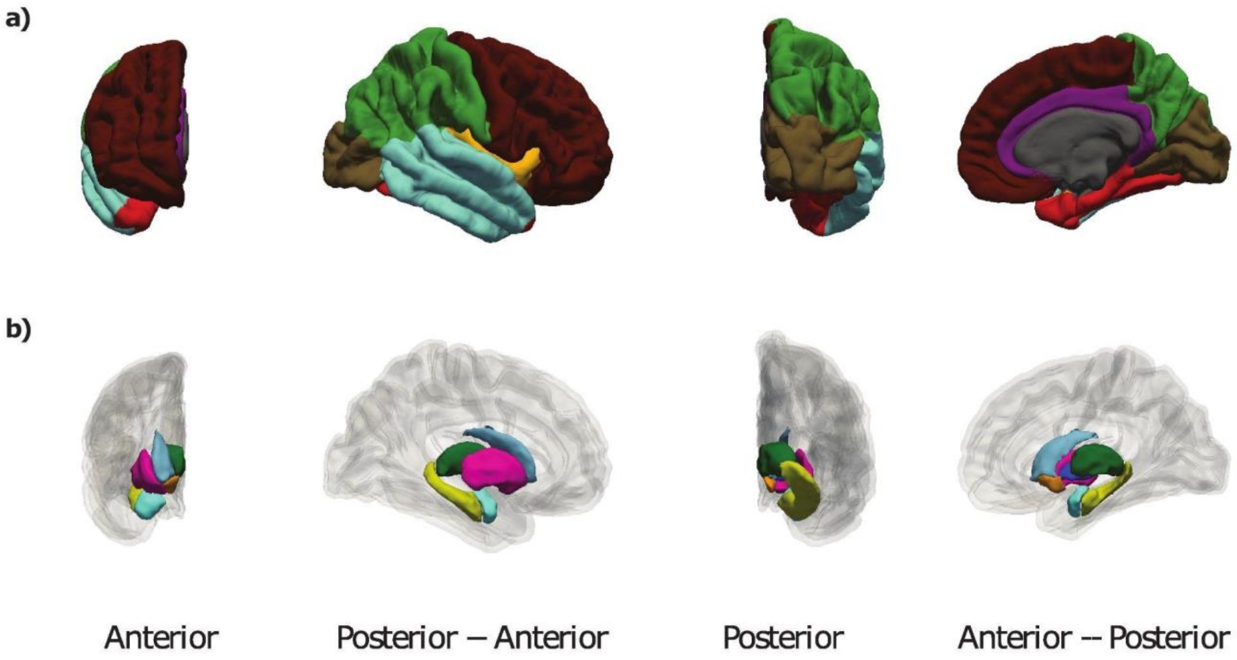


**Figure S1.1:** Panel **a)** cortical ROIs; Panel **b)** subcortical ROIs.

**Table S1:** The selected ROIs and their FreeSurfer label and Index. Sub-cortical regions are from the Aseg atlas and cortical regions from the aparc atlas.

| Region | ROI | Label | Index Left | Index Right |
|---|---|---|---|---|
| Sub-Cortical | Caudate | Caudate | 11 | 50 |
| | Accumbens | Accumbens | 26 | 58 |
| | Pallidum | Pallidum | 13 | 52 |
| | Putamen | Putamen | 12 | 51 |
| | Thalamus | Thalamus | 10 | 49 |
| | Hippocampus | Hippocampus | 17 | 53 |
| | Amygdala | Amygdala | 18 | 54 |
| Cortical | Insula | insula | 1035 | 2035 |
| | Frontal | paracentral | 1017 | 2017 |
| | | precentral | 1024 | 2024 |
| | | parsopercularis | 1018 | 2018 |
| | | parstriangularis | 1020 | 2020 |
| | | rostralmiddlefrontal | 1027 | 2027 |
| | | caudalmiddlefrontal | 1003 | 2003 |
| | | frontalpole | 1032 | 2032 |
| | | medialorbitofrontal | 1014 | 2014 |
| | | superiorfrontal | 1028 | 2028 |
| | | lateralorbitofrontal | 1012 | 2012 |
| | | parsorbitalis | 1019 | 2019 |
| | Temporal | inferiortemporal | 1009 | 2009 |
| | | middletemporal | 1015 | 2015 |
| | | bankssts | 1001 | 2001 |
| | | superiortemporal | 1030 | 2030 |
| | | transversetemporal | 1034 | 2034 |
| | Parietal | inferiorparietal | 1008 | 2008 |
| | | postcentral | 1022 | 2022 |
| | | superiorparietal | 1029 | 2029 |
| | | supramarginal | 1031 | 2031 |
| | | precuneus | 1025 | 2025 |
| | Occipital | lateraloccipital | 1011 | 2011 |
| | | cuneus | 1005 | 2005 |
| | | lingual | 1013 | 2013 |
| | | pericalcarine | 1021 | 2021 |
| | Parahippocampalgyrus | parahippocampal | 1016 | 2016 |
| | | entorhinal | 1006 | 2006 |
| | | temporalpole | 1033 | 2033 |
| | | fusiform | 1007 | 2007 |
| | Cingulate | caudalanteriorcingulate | 1002 | 2002 |
| | | rostralanteriorcingulate | 1026 | 2026 |
| | | isthmuscingulate | 1010 | 2010 |
| | | posteriorcingulate | 1023 | 2023 |

#### S1.2 Distribution Mapping (DiMap)

The distribution mapping (DiMap) approach (Properzi et al., 2019) was used to harmionize [11C]Cimbi-36 $BP_{ND}$ and [18F]altanserin $BP_P$. The basic idea is to estimate the underlying distribution and compute the Cummulative Density Function (CDF) for each tracer. Subsequently, both CDFs are used to derive a mapping between the two distributions, as visualized in **Fig S1.2**. The DiMap approach requires three assumptions to be fulfilled: 1 The rank order of the two measures is preserved, 2. Both measures are sampled from the same population and 3. Both tracers follow a specific distribution. Here, it was assumed that 1. and 2. hold true for the data used. Further, it was assumed that both the [18F]altanserim $BP_P$ and [11C]Cimbi36 $BP_{ND}$ values follow a normal distribution.

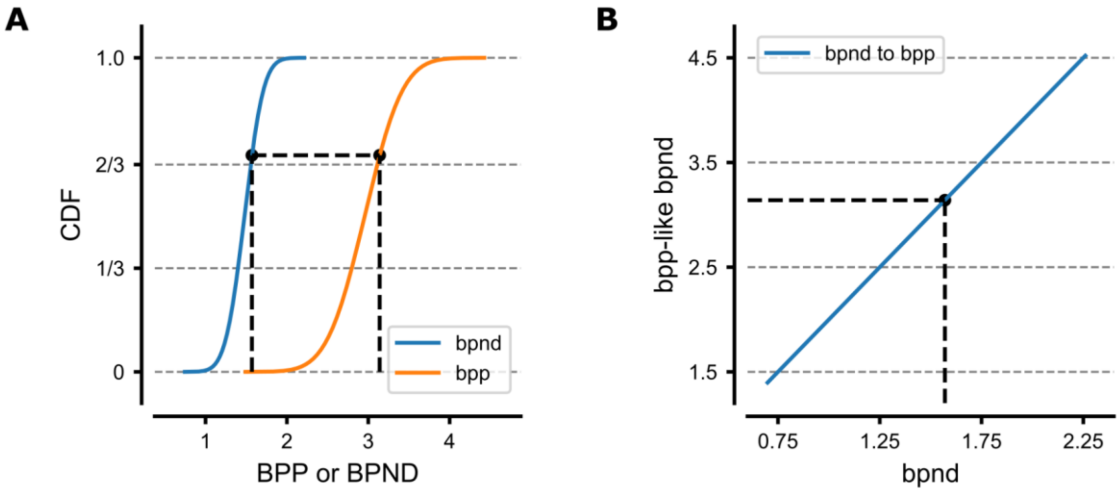


**Figure S1.2:** Illustration of Distribution mapping via the CDF. Panel **A**) visualizes two approximated CDFs for simulated data for two different samples illustrating $BP_{ND}$ and $BP_P$values. Panel **B**): shows the mapping from $BP_{ND}$ to $BP_P$ like $BP_{ND}$ values

### S2 Results

#### S2.1 Cross-Validation Results for all Models

**Table S2.1:** The results of all models for the four experiments (Reference, 5-HT2AR, GM, 5-HT2AR + GM). Values are mean (std) over the 100 CV iterations for MAE, R2, r age, and r pad.

| Modality | Model | E_gen (std) | R2<br>mean (std) | r_age<br>mean (std) | r_pad<br>mean (std) | p |
|---|---|---|---|---|---|---|
| Reference | pyment | 3.5 (0.4) | 0.93 (0.02) | 0.96 (0.01) | -0.36 (0.1) | <<0.01 |
| Reference | dummy | 13.33 (0.91) | -0.01 (0.01) | -0.0 (0.0) | -1.0 (0.0) | <<0.01 |
| 5-HT2AR | bridge | 6.63 (0.77) | 0.74 (0.07) | 0.87 (0.04) | -0.48 (0.15) | – |
| 5-HT2AR | rvm | 6.71 (0.78) | 0.73 (0.07) | 0.86 (0.04) | -0.47 (0.15) | 0,6 |
| 5-HT2AR | lingpr | 6.7 (0.8) | 0.73 (0.07) | 0.86 (0.04) | -0.49 (0.15) | 0,62 |
| 5-HT2AR | lsvr | 6.81 (0.77) | 0.72 (0.07) | 0.86 (0.04) | -0.49 (0.15) | 0,37 |
| 5-HT2AR | rbfgpr | 6.82 (0.8) | 0.68 (0.1) | 0.84 (0.06) | -0.62 (0.14) | 0,58 |
| GM | bridge | 7.2 (0.84) | 0.67 (0.13) | 0.84 (0.06) | -0.55 (0.16) | 0,3 |
| GM | rvm | 7.09 (0.86) | 0.68 (0.12) | 0.84 (0.06) | -0.53 (0.16) | 0,42 |
| GM | lingpr | 7.25 (0.83) | 0.67 (0.12) | 0.84 (0.06) | -0.55 (0.16) | 0,27 |
| GM | lsvr | 7.33 (0.88) | 0.66 (0.14) | 0.84 (0.05) | -0.49 (0.16) | 0,23 |
| GM | rbfgpr | 6.95 (0.83) | 0.69 (0.09) | 0.84 (0.05) | -0.61 (0.12) | 0,51 |
| 5-HT2AR + GM | bridge | 5.54 (0.7) | 0.8 (0.07) | 0.91 (0.03) | -0.37 (0.18) | 0,01 |
| 5-HT2AR + GM | rvm | 5.54 (0.68) | 0.8 (0.07) | 0.91 (0.03) | -0.39 (0.17) | 0,01 |
| 5-HT2AR + GM | lingpr | 5.6 (0.69) | 0.8 (0.07) | 0.9 (0.03) | -0.37 (0.17) | 0,01 |
| 5-HT2AR + GM | lsvr | 5.96 (0.68) | 0.78 (0.07) | 0.9 (0.03) | -0.39 (0.19) | 0,08 |
| 5-HT2AR + GM | rbfgpr | 5.69 (0.82) | 0.78 (0.09) | 0.9 (0.03) | -0.27 (0.17) | 0,07 |

r_age - Pearson's r between predicted and chronological age; r_pad - Pearson's r beteween PAD and chronological age; pl , pr - probabilities left and right of null-hypothesis in Bayesian setting. The null hypothesis is that there is no difference in MAE between the respective model and the base model across CV folds. Left: the base model has a lower MAE, right: the base has a higher MAE;

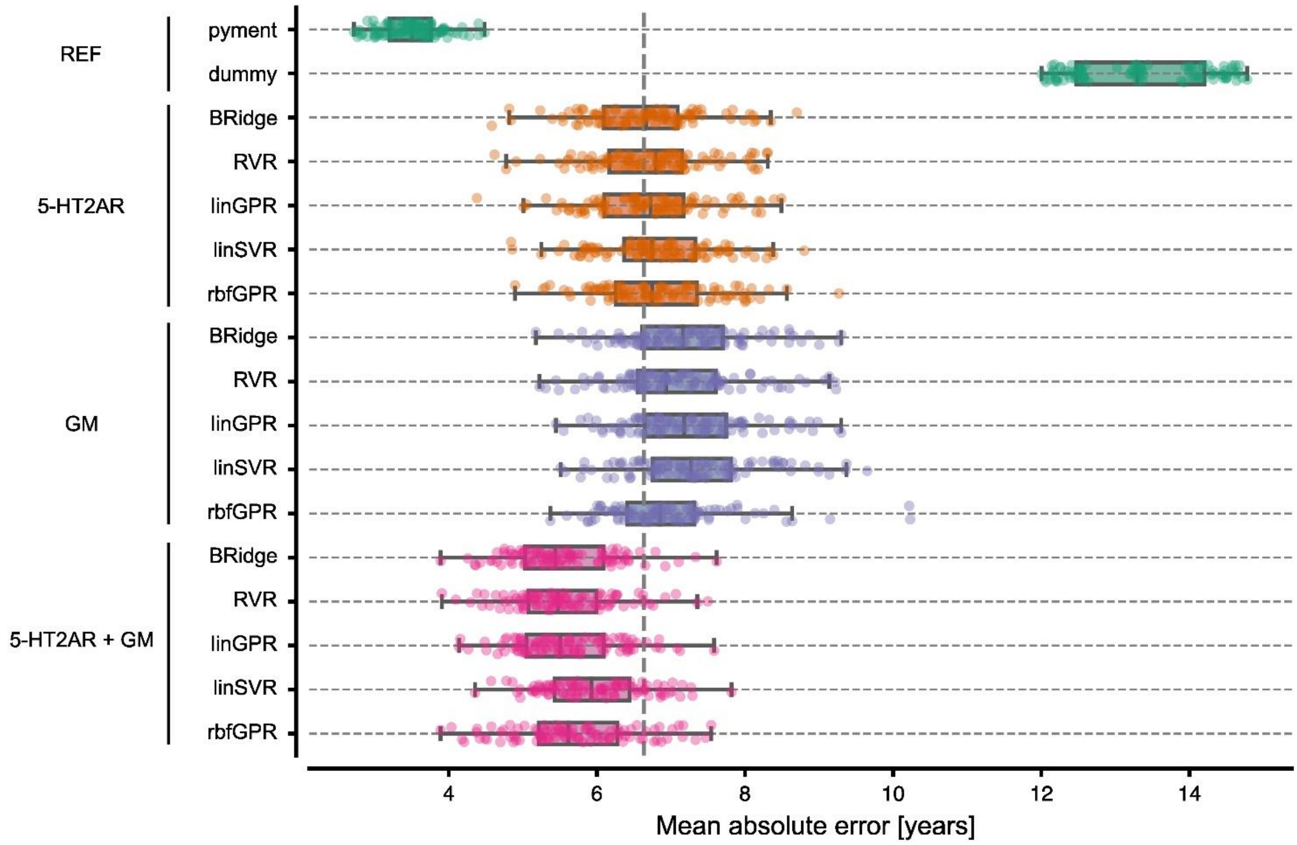


**Figure S2.1:** MAE for the different models. Each dot represents the mean MAE for one of the 100 CV iterations. The vertical dashed line marks the generalization error of the 5-HT2AR binding based model.

#### S2.2 Feature Weights

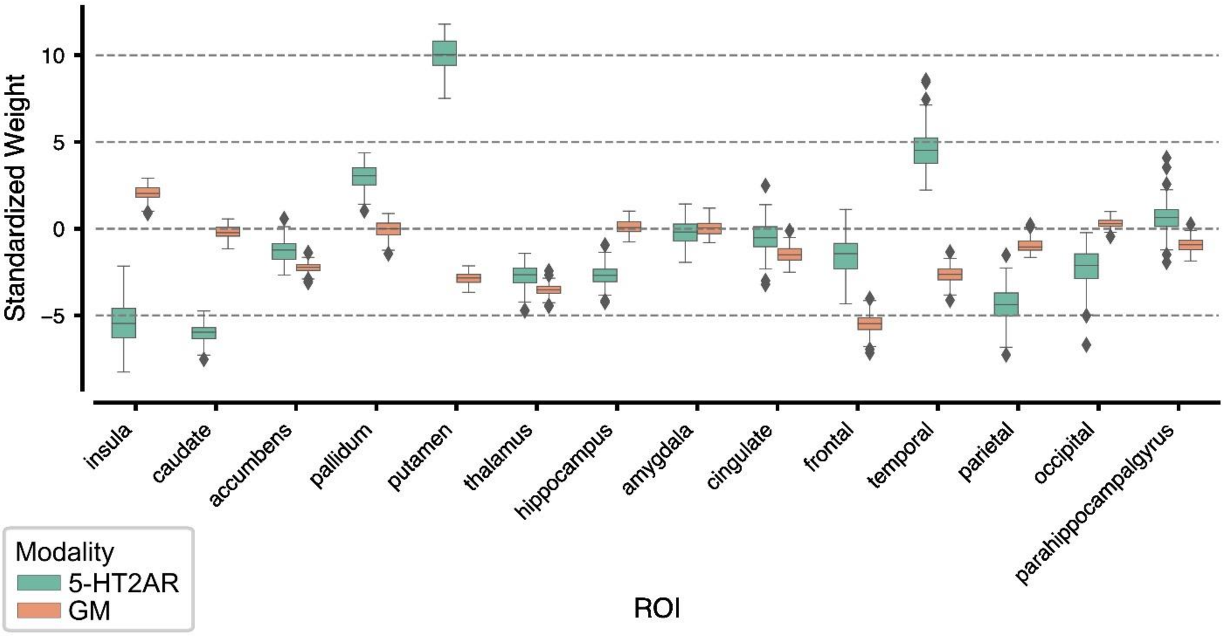


**Figure S2.2:** Feature weights for each modality in the stacking regressor using BRidge as base algorithm. The whiskers indicate 1.5 times the interquartile range. The boxplots represent the distribution of the weights for each of the 100 CV folds.
